# Echoes of the late Pleistocene in a novel trophic cascade between cougars and feral donkeys

**DOI:** 10.1101/2021.04.13.439662

**Authors:** Erick J. Lundgren, Daniel Ramp, Owen M. Middleton, Mairin Balisi, William J. Ripple, Chris D. Hasselerharm, Jessica N. Sanchez, Eamonn I. F. Wooster, Mystyn Mills, Arian D. Wallach

## Abstract

Introduced large herbivores have partly filled ecological gaps formed in the late Pleistocene, when many of the Earth’s megafauna were driven extinct. However, surviving predators are widely considered unable to influence introduced megafauna, leading them to exert unusually strong herbivory and disturbance-related effects. We report on a behaviorally-mediated trophic cascade between cougars (*Puma concolor*) and feral donkeys (*Equus africanus asinus*) at desert wetlands in North America. In response to predation of juveniles, donkeys shifted from nocturnal to almost exclusively diurnal, thereby avoiding peaks in cougar activity. Furthermore, donkeys reduced the time they spent at desert wetlands by 87%: from 5.5 hours a day to 0.7 hours at sites with predation. These shifts in activity were associated with increased activity and richness of other mammal species and reduced disturbance and herbivory-related effects on these ecologically-distinct wetland ecosystems, including 49% fewer trails, 35% less trampled bare ground, and 227% more canopy cover. Cougar predation on introduced donkeys rewires an ancient food web, with diverse implications for modern ecosystems.

## Introduction

Many of the world’s largest herbivores and predators were lost in the late Pleistocene, most likely due to human hunting [1, 2]. A second wave of declines is ongoing, as the majority of surviving large herbivore species (henceforth megafauna) are now threatened with extinction [3]. Yet several megafauna species have also been introduced, thereby restoring lost species richness and potential ecological functions [4, 5]. However, it has long been assumed that surviving predators are incapable of exerting ecologically-significant predation pressure on introduced megafauna, leading to unusually strong herbivory and disturbance-related effects on modern ecosystems relative to native megafauna (Fig. 1a-b).

**Fig 1.**
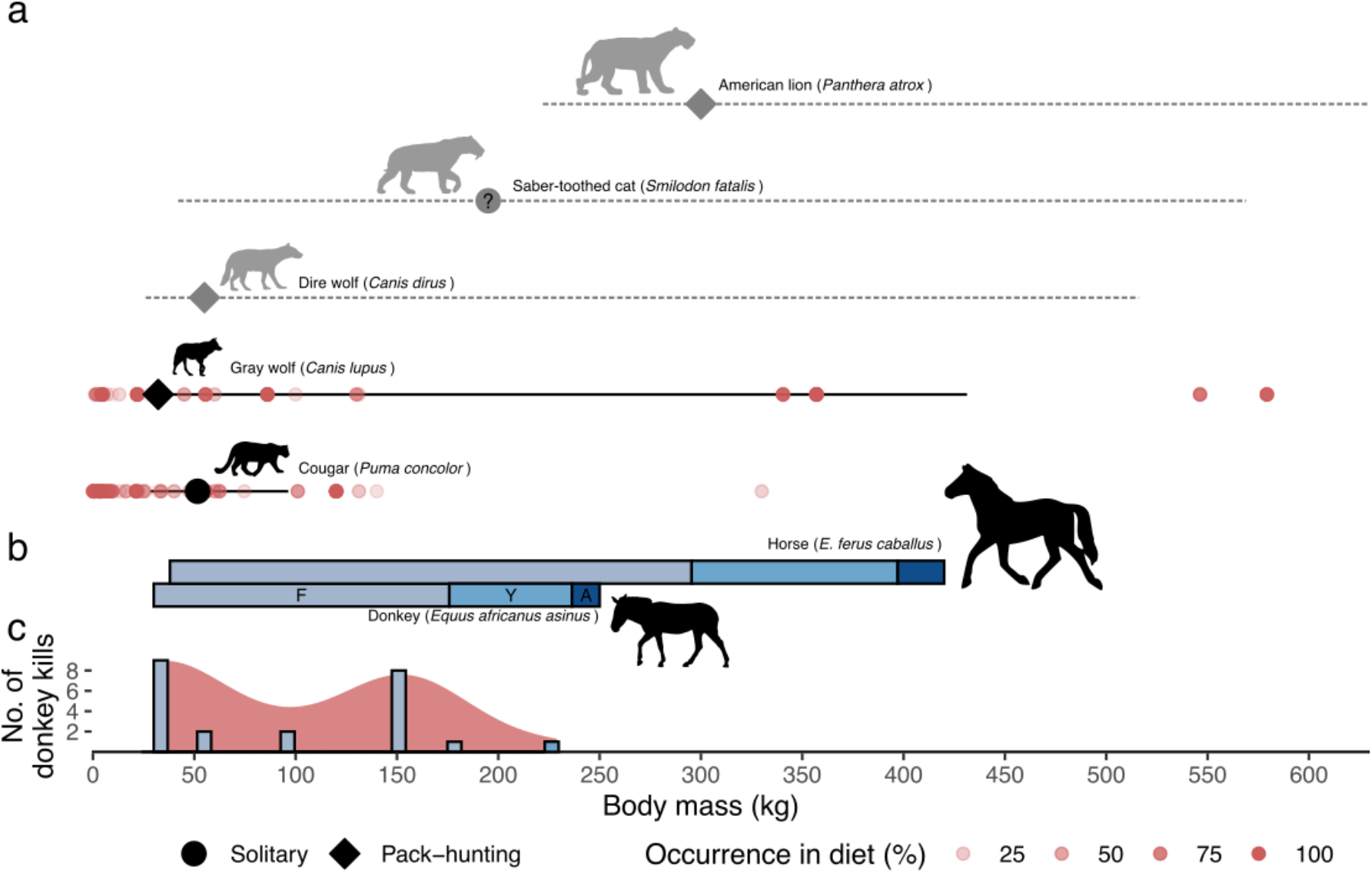
Body size and hunting style determine predator-prey interactions and may constrain the ability of extant predators to influence introduced megafauna. X-axis (body mass) is shared across all subplots. **a.** North American apex predators before and after the late Pleistocene extinctions. Horizontal lines indicate theoretical optimum prey body mass range of extant (black) and extinct (dashed) predators [from 12]. Points indicate average predator body mass and hunting style, which remains uncertain for *Smilodon fatalis* (denoted with question mark). Red points indicate published prey items by body mass, with transparency denoting the frequency of prey occurrence in diet [data from 33]. Of extant predators, only the cougar (*Puma concolor*) substantially overlaps in geographic distribution with introduced equids in North America [29, 51]. **b.** Estimated body mass ranges for equid age classes (F=foal, Y=yearling, A=adult). **c.** Body mass density distribution (red fill) and frequency (overlaid bars) of cougar donkey kills from field surveys (this study).

We report on a novel trophic cascade [6] between cougars (*Puma concolor*) and feral donkeys (*Equus africanus asinus*) in North America. Cougars co-occurred with a diversity of equid species for more than a million years until the North and South American late Pleistocene extinctions ~9-12,000 years ago [7]. Paleoecological evidence, however, suggests that cougar-equid interactions may have been uncommon, with equids mainly preyed upon by larger or pack-hunting now-extinct predators [Fig. 1a-b, 8]. Cougars are likewise widely considered in research and policy as incapable of significantly influencing introduced equid populations or ecologies [9]. Despite this, we mapped widespread predation of juvenile donkeys across the Sonoran and Mojave Deserts of North America (Fig. 1c) and recorded the first, to our knowledge, documented predation of a yearling (Fig. 2a-b) and foal (Fig. 2c-d).

**Fig 2.**
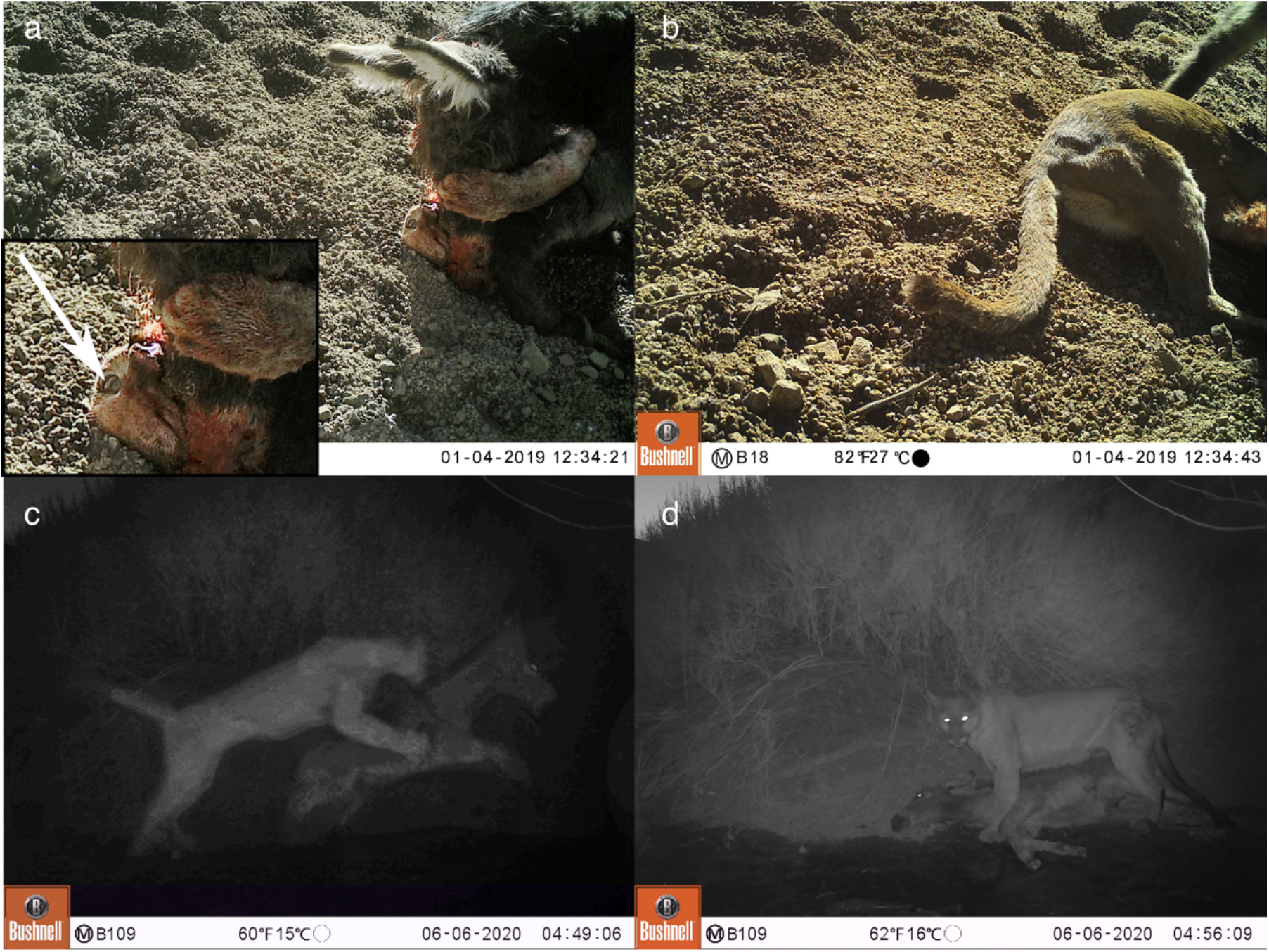
First reported predation of feral donkeys by cougars, captured by trail camera. **a-b.** Successful predation of a yearling donkey in the Sonoran Desert, Arizona. The cougar is looking up from the ground. Arrow in inset points to the cougar’s left eye. **c-d.** Predation of a foal in Death Valley National Park, Mojave Desert, California. Donkey ages were determined from tooth eruption sequences of carcasses. Images **a** and **c** were tonally adjusted for visibility (see S1 Fig for original versions).

While predation influences populations through direct killing, it also drives predator-avoidance behaviors with cascading implications for the effects of herbivores on ecosystems [10]. However, it remains contested whether predation of juveniles can drive ecologically-relevant, behaviorally-mediated trophic cascades, as invulnerable adults may not respond to predation risk [11–13]. To answer this question, we investigated whether predation by cougars influenced donkey behavior and their associated effects on other vertebrates and vegetation at desert wetlands.

## Results

To understand if donkeys altered their behavior to avoid cougar predation, we compiled trail camera images from 24 desert wetlands in the Sonoran and Mojave Deserts spanning 9,303 trap nights (S2 Fig, S1 Table). We compared donkey activity patterns between wetlands in regions with low densities of cougars and no evidence of predation (henceforth ‘low predation risk’) and regions with abundant cougars and widespread predation (‘high predation risk’). The high predation risk regions also included sites shielded by human recreational activity where cougars were locally absent, which were treated separately (‘human shielding’ [14]).

Donkeys significantly changed their behavior in response to predation risk (Fig 3a, S3 Fig). Donkeys were primarily nocturnal at low predation risk wetlands, but became almost exclusively diurnal at high predation risk wetlands, thereby avoiding the peak of cougar activity (Fig 3a). Indeed, the nighttime attack captured on camera (Fig 2c-d) was one of the few nighttime donkey visits to that wetland–with immediate consequences for the foal. Corroborating that these behavioral changes were a direct response to predation risk from cougars, donkeys were again nocturnal at human-shielded sites within this same high predation risk region (Fig 3a).

**Fig 3.**
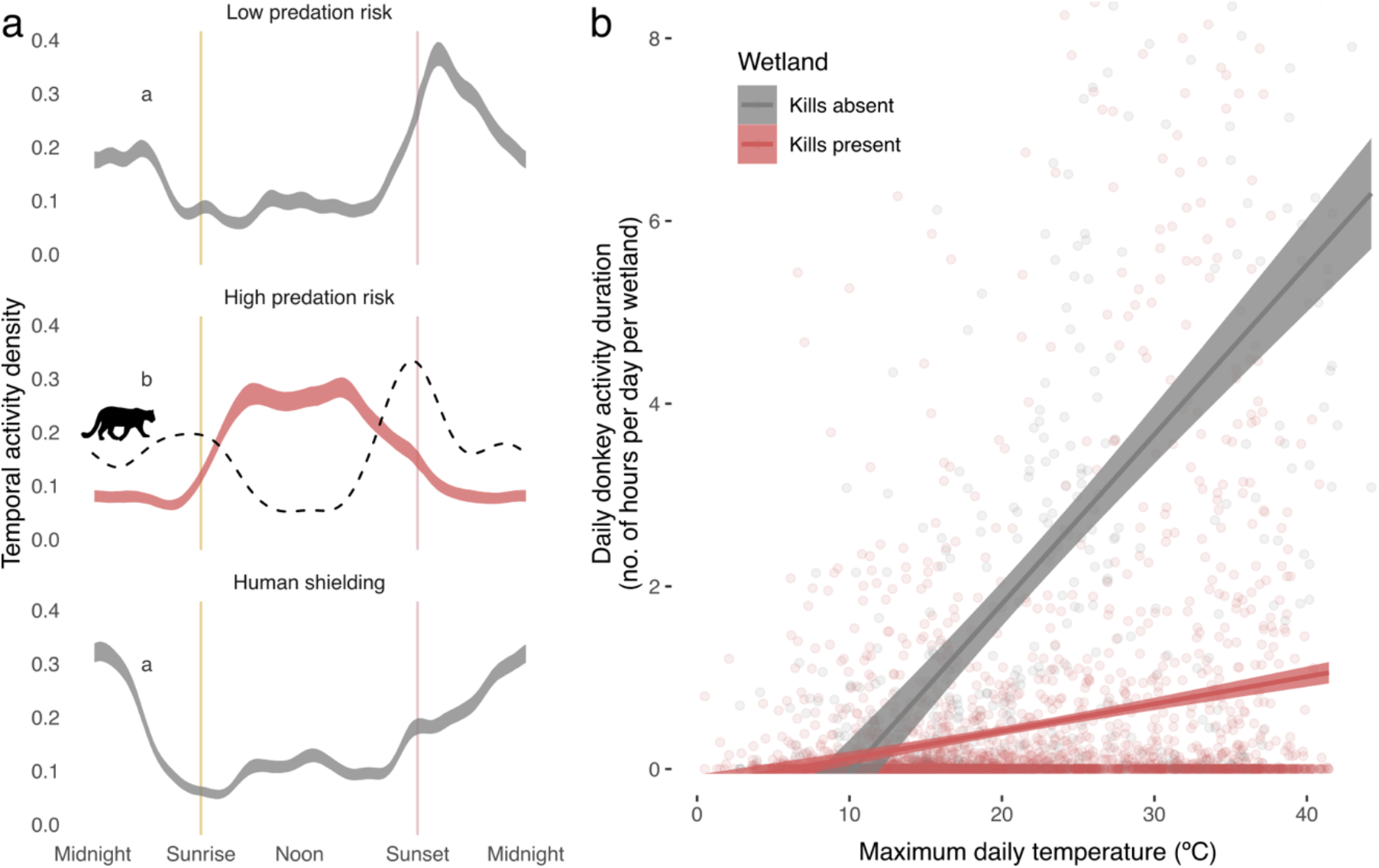
Cougars shape donkey behavior at desert wetlands. **a.** Donkey temporal activity under different levels of predation risk. X-axis indicates time of day. Ribbon indicates donkey activity 95% confidence intervals from bootstrapped detections, with letters indicating significance groupings based on non-overlap of confidence intervals (see S3 Fig). Dashed line indicates cougar activity pattern. ‘Low predation risk’ = sites in regions with low densities of cougars where predation was absent, ‘high predation risk’ = sites in regions with cougars and widespread predation, ‘human shielding’ = wetlands with recreational activity where cougars were absent, despite proximity to high predation risk wetlands. **b.** Relationship between maximum daily temperature and the activity of donkeys at wetlands (hours/day/site) for sites with and without predation.

Temperature drives water requirements, particularly in desert environments [15]. Desert wetlands can thus become foci of activity for water-dependent animals such as donkeys during the summer. To understand if cougars mediate this interaction, we calculated the daily activity of donkeys at each wetland (hours/day/site) from trail camera data and analyzed how it was shaped by maximum daily temperature and two indicators of predation risk: the presence of kills (cached carcasses) and the presence of cougars. We used multimodel inference (based on AICc) to choose the most parsimonious model.

Donkey activity was affected by the presence of kills (*χ*^*2*^=300.4, p<0.001), maximum daily temperature (*χ*^*2*^=55.4, p<0.001), and their interaction (*χ*^*2*^=14.6, p=0.001). At wetlands without kills, activity reached an average of 5.5 hours/day on days ≥35°C (SD=±4.4, max=16.5, Fig 3b). However, at sites with kills, activity remained low and relatively stable regardless of temperature, averaging 0.7 hours/day on days ≥35°C (±1.7, max=12.6, Fig 3b). The presence of cougars (i.e., at sites with cougars but no kills) also affected activity, but to a lesser extent and was not included in the most parsimonious model (S4 Fig).

Megafauna, including introduced equids, can competitively exclude smaller species from water points [16, 17]. We thus identified all mammal species larger than a cottontail rabbit (*Sylvilagus audubonii*) from camera trap imagery and assessed whether cougar predation on donkeys, and corresponding changes in donkey behavior, was associated with increased use of wetlands by other species.

As with donkeys, the activity duration of other mammals increased with maximum daily temperature (Fig 4a, z=8.1, p<0.001). However, this was strongest at wetlands with cougar predation, where donkey activity was reduced (Fig 4a, post hoc t-ratio=−6.46, p<0.001). Indeed, across all sites, the daily activity of other mammals was negatively related to the activity of donkeys on that same day (Fig 4b z=4.5, p<0.001), which further supports that cougar predation on donkeys itself, not other factors, facilitates the activity of other mammal species at these wetlands. These increases in activity corresponded with higher mammal species richness at sites with active donkey predation (Fig 4c, W=37.5, p=.045).

**Fig 4.**
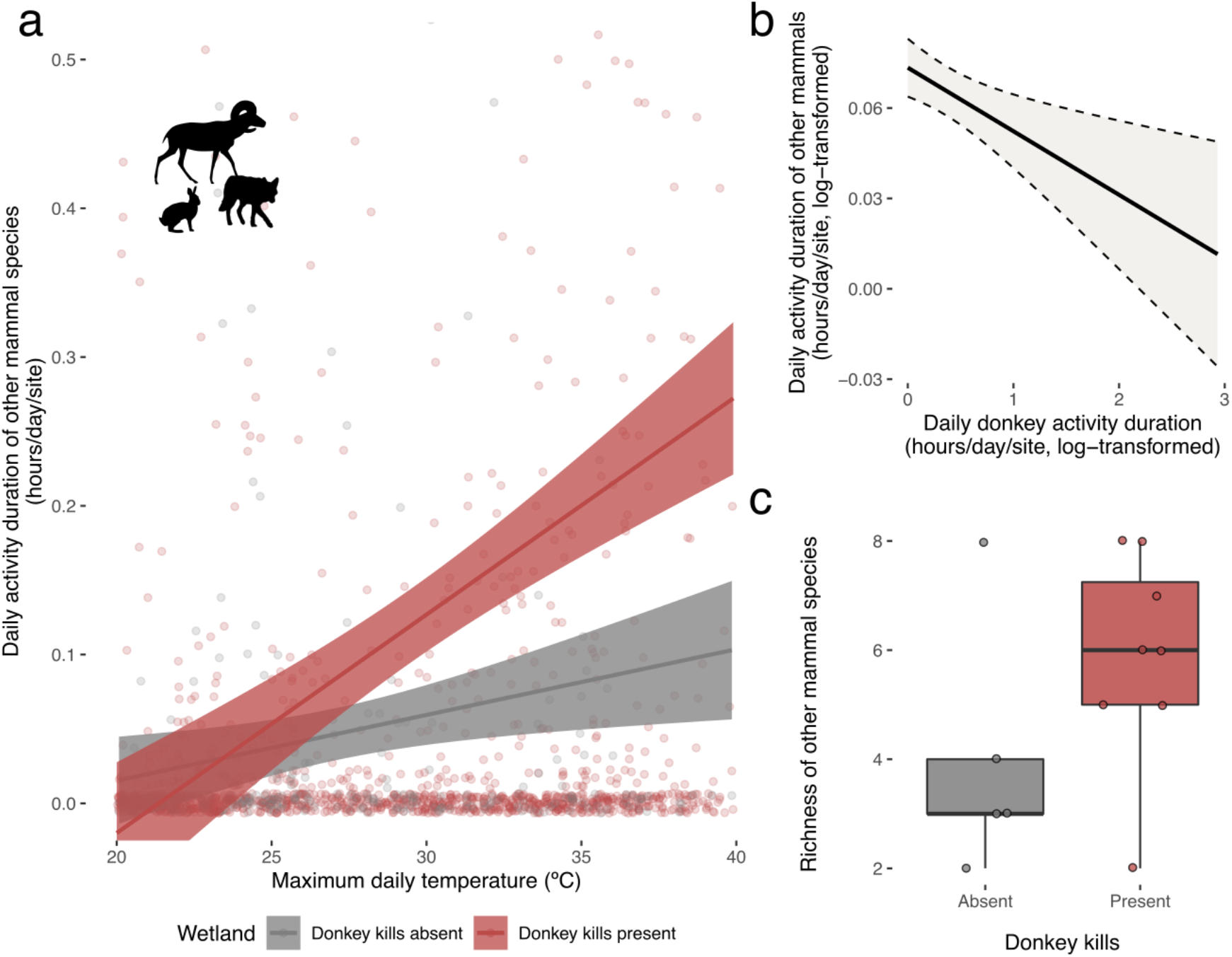
Predation on feral donkeys was associated with increased utilization of wetlands by other mammal species. **a.** Duration of activity by other mammal species at wetlands increased with maximum daily temperature (*χ*^*2*^=210.25, p<0.001), but to a greater extent at sites with active donkey predation (interaction term: t-ratio=−6.42, p<0.001). **b.** Daily activity duration of other mammals had a negative relationship to the activity of donkeys on that day, across all sites (z=4.5, p<0.001). Durations were log-transformed to reduce over dispersion. **c.** Richness of other mammal species at sites with and without donkey kills (W=37.5, p=.045). Given unequal trap nights, richness was interpolated following [52].

**Fig 5.**
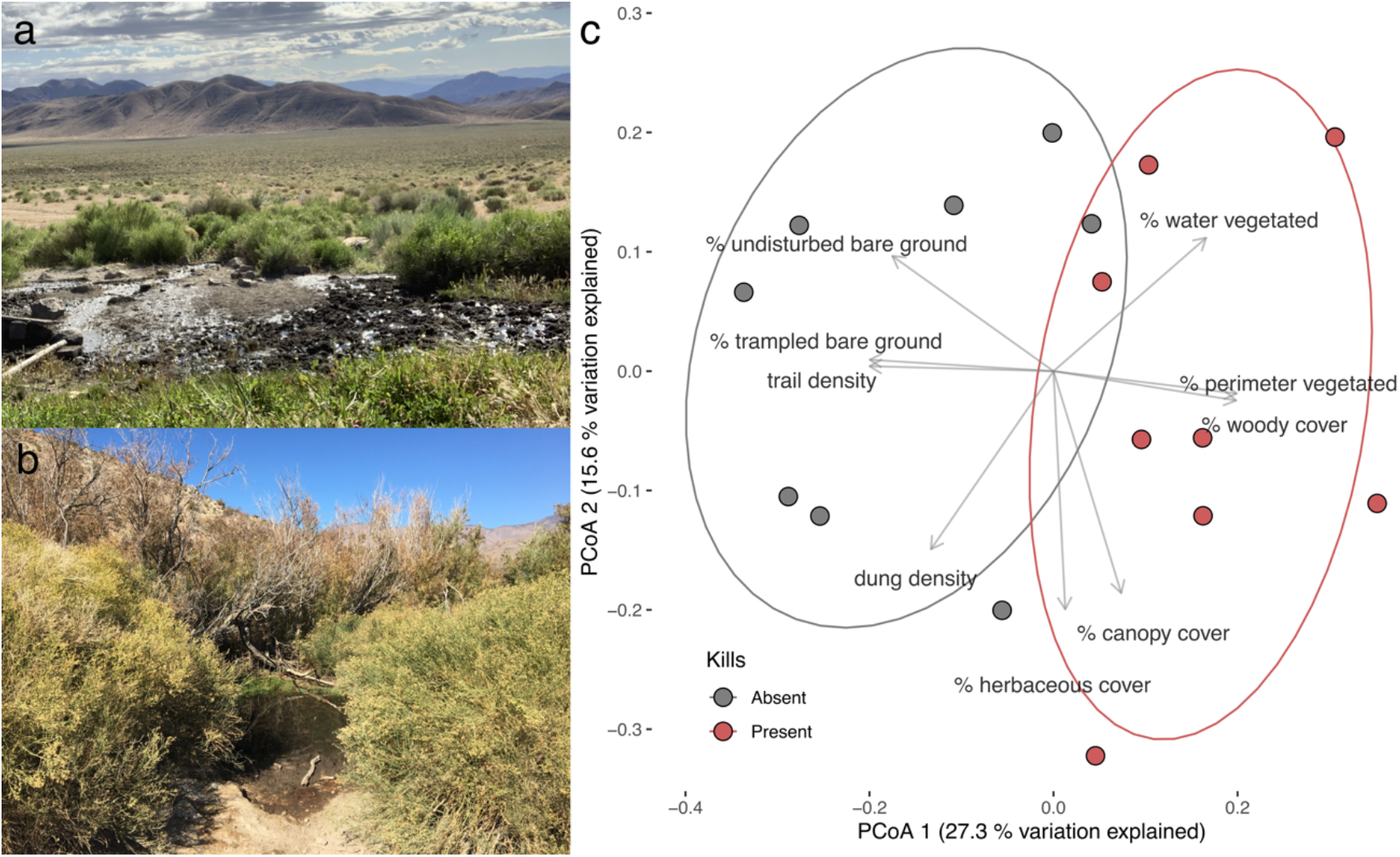
Cougar predation is associated with reduced herbivory and disturbance-related influences on desert wetlands. **A.** A representative wetland lacking both cougars and kills compared to a nearby (~6km) wetland (**B**) where cougars and kills were present (site of the kill in Fig 2C-D). Photos were taken at a similar distance from water’s edge, by EJL (**A**) and OMM (**B**). **C.** Principal coordinates analysis (PCoA) showing significant differences in wetland structure with and without cougar predation on donkeys (PERMANOVA, F=3.5, p<0.001). Each point is a wetland. Relationship between response variables and PCoA axes are indicated by overlaid arrows and text. The presence of cougars themselves, elevation, and terrain complexity were not significant (PERMANOVA p=0.27-0.33), nor was the geographic distances between sites (F=0.82, p=0.58). See S5 Fig for response of individual variables.

Cougar predation on donkeys was also associated with reduced herbivory and disturbance-related effects on these ecosystems. We collected data on nine soil and vegetation responses encompassing potential disturbance and herbivory-related effects of donkeys on desert wetlands (S6 Fig). We synthesized these data with a Principal Coordinates Analysis (PCoA) to find the primary axes by which wetlands differed from each other, which revealed significant differences between sites with and without kills (PERMANOVA: R^2^=0.20, F=3.54, p<0.001, Fig 4a-c). Wetlands with kills had more vegetation, including 227% more canopy cover (from 12.7±16% to 42±34%, mean±SD, S5 Fig), and 183% more vegetation around water perimeter (from 21±21% to 60±18%, S5 Fig). Likewise, these sites had less disturbance, including ~49% fewer trails to water (from 3.2±2, to 1.6±0.7, S5 Fig) and 35% less trampled bare ground (from 77±20% to 50%±16%, S5 Fig).

The presence of cougars themselves (independent of kills), topographic complexity, and elevation did not explain dissimilarity in wetland structure (p=0.27−0.33, S2 Table). Likewise, geographic distances between sites did not influence wetland dissimilarity (multiple regression on distance matrices, R^2^=0.007, F=0.82, p=0.58), indicating that the differences between wetlands were driven by predation upon donkeys and not by underlying spatial gradients.

## Discussion

For more than a million years, equids co-occurred with cougars and much larger predators, of which the latter were likely their primary predators [8]. Cougar predation on juvenile donkeys nonetheless was strongly associated with altered donkey behavior, increased activity and richness of other mammals, and reduced effects on wetland vegetation and soil. This adds to growing evidence that extant predators have a greater capacity to influence introduced equids than typically considered. This includes wolves (*Canis lupus*) and brown bears (*Ursus arctos*) in Eurasia and North America, jaguars (*Panthera onca*) in North and South America, and dingoes (*Canis dingo*) in Australia (Fig 6, S3 Table). Cougars can be significant predators of introduced horses as well [18, 19], suggesting the possibility of behaviorally-mediated trophic cascades among these larger megafauna [6] (Fig 6). The return of equids after their ~12,000 year hiatus–and predation upon them by cougars–suggests a rewiring of food webs [20], with diverse implications for modern ecosystems, the cougars and their prey, and for how these species are treated in policy.

**Fig 6.**
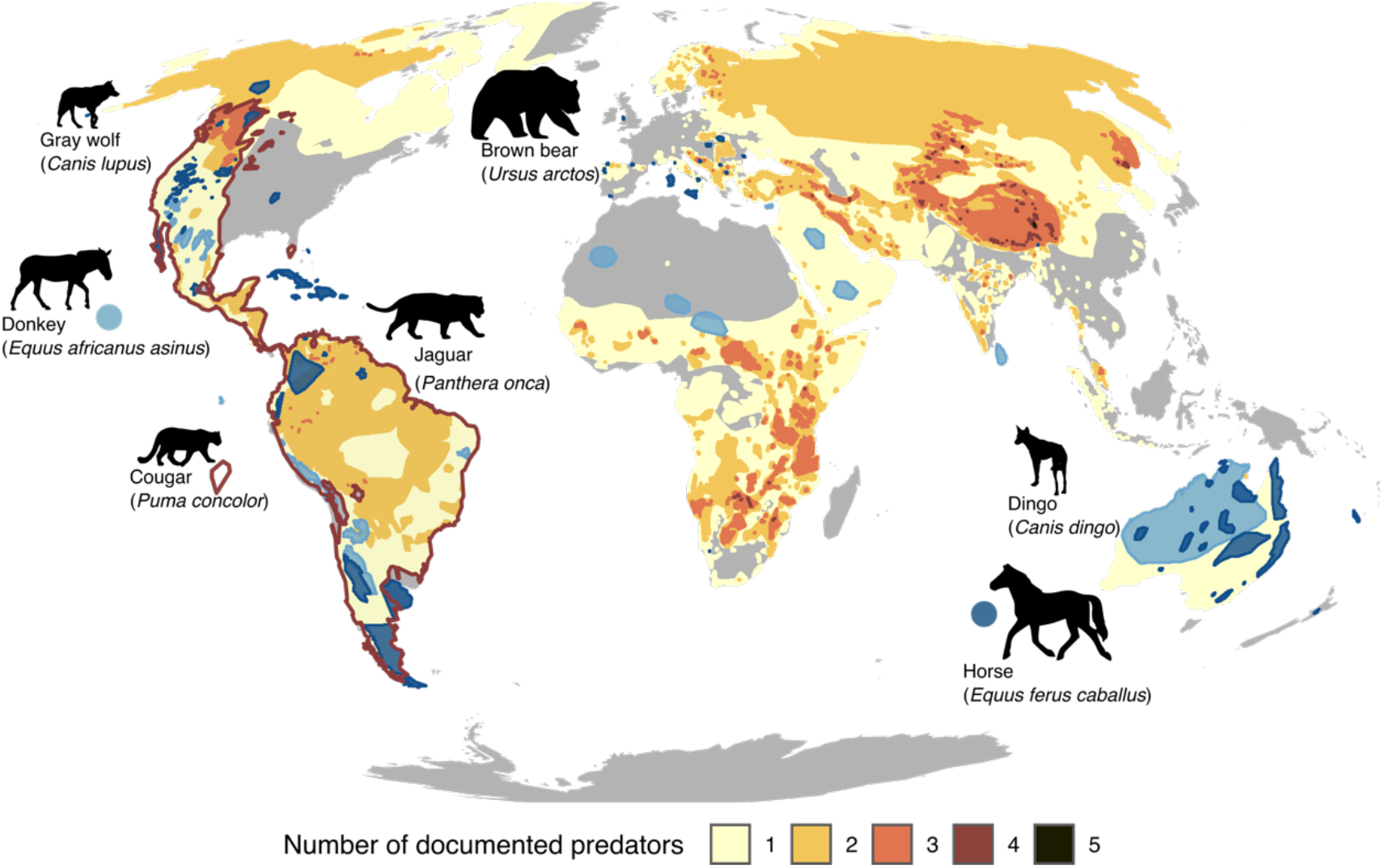
Distribution of introduced equids and species richness of predators documented predating on equids. Introduced ranges of horses (dark blue) and donkeys (light blue) are overlaid on predator species richness (yellow-orange-black gradient). Cougar distribution is demarcated with red border overlay. The predators included have all been documented preying on equids. Those highlighted with icons (see **S3 Table**) have the potential to overlap in distribution with feral equids. Feral equids are among the most widespread and abundant introduced megafauna species **[5, 29]**. Though rarely studied, the predators that survived the late Pleistocene extinctions have greater capacity to influence feral equids than usually considered. Predator range maps were drawn from the IUCN Redlist **[51]**, except for the dingo **[53]**. Introduced equid ranges are from **[29]**.

The influences of mammalian megafauna on wetlands, particularly in water-limited drylands, were likely ubiquitous from the early Cenozoic (30-40 million ybp) until the late Pleistocene extinctions [1, 4]. Megafauna well digging, disturbance, and herbivory can maintain wetland heterogeneity and water availability [21, 22]. However, the effects of prehistoric megafauna, even those ≥1,000 kg, were likely mediated by predation, if only on juveniles [12]–potentially an ancient parallel to the behaviorally-mediated trophic cascade driven by cougar predation on juvenile donkeys.

Donkeys, cougars, and smaller-bodied prey of cougars, such as bighorn sheep (*Ovis canadensis*) and mule deer (*Odocoileus hemionus*), are entangled in an emerging ecological network. In addition to shaping donkey ecology, cougar predation on donkeys may thus engender novel evolutionary trajectories in cougars [23] and reduce pressure on their other prey. These relationships may also yield overlooked consequences in response to ongoing donkey removals and near-ubiquitous cougar persecution. Donkeys were the primary recorded prey of cougars at our study sites (24 of 29 cached carcasses) and horses have been documented as primary prey elsewhere [19]: removing equids may thus lead to increased predation on smaller prey [24].

Likewise, killing cougars in service of the livestock industry and to increase populations of deer and bighorn sheep for conservation and sport hunting [25, 26] may influence feral equid populations and their ecological effects. As with many apex predators, persecution can reduce the ability of cougar populations to hunt larger, more challenging prey by removing older individuals and by disrupting the transmission of hunting techniques from mother to young [27]. Even moderate persecution may thus reduce the potential for ecologically-significant trophic cascades between cougars and feral equids, despite their broad geographic overlap (Fig 6).

Cougars avoided human recreation sites and were constrained by their need for topographic or vegetative ambush cover to hunt successfully (S7 Fig) [28]. This created a mosaic of predation risk, which our study design utilized. However, without experimental manipulation of cougar presence, the stark differences observed between wetlands with and without predation remain correlative. Despite this, we argue that the proximity (e.g., ~2-6 km) of wetlands shielded by recreation, where vegetative cover was non-existent, to topographically similar wetlands with active predation and abundant vegetation, indicates that cougar predation was the primary driver of observed differences in mammal richness and activity and wetland structure.

Horses and donkeys, like the majority of extant megafauna, are threatened in their native ranges [5]. This has led to calls for more inclusive conservation approaches [29], which may find productive common ground with conservation efforts to increase protection and tolerance for cougars and other predators. Doing so may further influence the ecologies of feral donkeys. For instance, the effects of human shielding and the ambush cover requirements of cougars suggest that managing recreation and allowing for the reestablishment of pack-hunting wolves (*Canis lupus*) or larger-bodied jaguars (*Panthera onca*) would shift predation risk into places currently deemed safe, with broad implications for how feral equids influence ecosystems.

The effects of organisms emerge in relation to ecological contexts and are not essential to their dispersal histories, whether of unassisted colonization or human introduction [30–32]. If we had conducted this research to enumerate the effects of feral donkeys on desert wetlands, without quantifying cougar predation, our data would contain a great degree of inexplicable noise. Instead we find echoes of the late Pleistocene in a novel trophic cascade between cougars and feral donkeys, with diverse ecological and evolutionary consequences.

## Materials and Methods

### Theoretical predator prey ranges

We followed Van Valkenburgh et al. 2016’s [12] formulae to calculate theoretical optimal prey sizes based on allometric scaling relationships from observed diets in extant predators (Fig 1). Pack-hunting and solitary hunting (e.g., cougars) species were treated with separate equations, given the ability of pack-hunting species to cooperatively kill larger prey [12].

Importance of observed prey items for cougars and gray wolves were derived from CarniDIET [33]. We only included studies that reported the frequency of a prey item across predator scats in a study. Predator and prey body masses were derived from Mass of Mammals [34]. Donkey and horse body masses by age class were estimated from scaling formulae in [35] and were calculated from adult body masses and birth weights reported in AnAge: The Animal Ageing and Longevity Database (36).

### Carcass and cougar surveys

We surveyed for cougar presence and donkey carcasses at 27 wetlands in Death Valley National Park. Surveys were conducted by walking the perimeter of wetlands and investigating every potential trail for cached carcasses [37]. Carcasses and cougar sign were frequently encountered on tight <0.5 m tall trails through dense riparian vegetation. Cougar scats were identified visually by size and shape and their frequent association with very large scrapes. All surveys and scat identification were conducted by EJL, thus eliminating inter-observer variability.

We classified donkey remains as kills if they were located on cougar trails or if they were within 20 m of a cougar scat or scrape. Cougar trails usually terminated at the kill and were defined by their height (<1 m) and because they were unused by living donkeys–as determined by the lack of donkey track and sign and by camera traps. We estimated donkey ages by examining tooth wear and eruption sequences following resources for donkeys and horses [38, 39]. In some cases, where skulls could not be located, we estimated age by the fusion or lack thereof of appendicular bones. We estimated the ages of the two photographically captured kills (Fig 2) by locating the carcasses and evaluating tooth eruption sequences as well.

### Activity patterns of donkeys and cougars

We collected ca. 2.5 million trail camera images (Bushnell Trail Cam Pro) from over five years and 64 camera stations across 26 wetlands in the Sonoran and Mojave Deserts of North America (S3 Fig, S1 Table). Sites averaged 388 trap nights per site, with a total of 9,303 trap nights across all sites and all seasons (S1 Table). Sites with less than 10 trap nights (*n* = 2) were excluded from analysis. All trail cameras were on water or on trails to water. Some sites contained multiple cameras, depending on the number of water-access points at each site, which were aggregated for analysis.

We evaluated the effect of cougars on the temporal patterns of feral donkeys with the overlap (v0.3.3) and circular packages (v0.4-93) in R (v4.0.0). To do so, we converted clock time to sun time, which is consistent relative to sunrise and sunset and is derived from the date and geographic coordinates of each trail camera [40, 41]. We compared donkey daily activity patterns between wetlands in regions with low densities of cougars and no evidence of predation (henceforth ‘low predation risk’) and regions with abundant cougars and widespread predation (‘high predation risk’). This latter region also included sites that were shielded by human activity, where cougars were locally absent [‘human shielding’, 14], despite proximity (e.g. <2 km) to wetlands with abundant cougars and predation.

Sampling effort (e.g., number of trap nights) and the number of donkey detections varied between sites and between predation risk categories, which would bias pooled estimates [42]. We therefore resampled our data over 1,000 bootstraps, sampling equally between predation-risk categories and sites within each category. We selected 25% of the number of detections in the predation-risk category with the lowest number of detections, and then drew this quantity equally from each predation-risk category, divided equally from all sites within each category. From each subset we calculated donkey temporal activity using a circular von Mises density distribution kernel, as appropriate for time data [41]. From these resampled density distributions, we calculated 95% confidence intervals (CIs) to test if there was a significant difference between activity patterns under different levels of predation-risk, based on the visual overlap or non-overlap of CIs.

To understand how cougars influenced the extent to which donkeys utilized wetlands, we calculated the daily activity duration of donkeys at each wetland by assigning donkey detections into events, defined as any detection ≥30 minutes apart from any other. We then summarized the total event duration per day and site. Given that study sites ranged across several distinct populations, including low density ones, we focused on one large contiguous population in the Southern Panamint Mountains of Death Valley National Park (permit number DEVA-2018-SCI-0036). To avoid potential bias, we monitored every identifiable wetland within this region, although camera thefts and malfunctions prevented the inclusion of some sites. Unfortunately, density estimates were not available. This population consisted of 15 wetlands, (3,746 trap nights) and included sites with kills (*n* = 9) and without (*n* = 6) and sites with camera trap detections of cougars (*n* = 9) and without (*n* = 6). Note that the presence of cougars and kills did not perfectly correspond: one site had cougar detections but no kills, another had clear evidence of kills within the last year, but no detections were made during the study period (S2 Fig, S1 Table).

Given that surface-water dependency is driven by temperature we extracted daily maximum temperatures from a 4×4 km interpolated national dataset for each site [43]. We then analyzed the effect of local kills, cougar presence, daily maximum temperature, and their interactions on daily activity with a negative binomial mixed effect model in the R package ‘glmmTMB’ v1.0.2.1 [44], nesting day within site as random effects. The presence of kills and cougars were based on study-wide presence or absence. We used multimodel inference (based on AICc) to remove spurious terms and selected the most parsimonious model, which included daily maximum temperature, local predation, and their interaction. Cougar presence itself was not included.

### Vertebrate activity patterns

To understand how donkey activity patterns may in turn alter the activity of other vertebrate species, we further analyzed trail camera imagery, focusing again on the Southern Panamints Mountains of Death Valley to control for differences in donkey population sizes between study areas. Given the large number of images, we only identified mammals larger than a cottontail rabbit (*Sylvilagus audubonii*). As with the donkey images, images of the same species were grouped into ‘events’ based on a 30-minute window.

We then analyzed how mammal activity (hours/day/site) varied by maximum daily temperature and the presence of kills, using a zero-inflated negative binomial model, nesting day within site. To better understand how this related to donkey activity, we analyzed how daily mammal activity related to the donkey activity of that same day, using a negative binomial distribution and nesting day within site as random effects. To assess if this further corresponded to differences in species richness, we used the R package ‘iNEXT’ v2.0.20, to interpolate richness (Hill order 0), accounting for differences in sampling effort at each site [45]. We tested for differences in richness between sites with and without donkey predation with a Wilcoxon signed rank test for non-parametric data.

### Effects of feral donkeys on wetlands

To understand if cougars influence the effects of donkeys on wetlands we collected data from 16 desert wetlands in Death Valley National Park in November of 2019. We focused on pools where donkeys accessed water, at which we measured the percent of surface water vegetated, the percent of water surface covered by canopy foliage, the number of access trails per pool, and the percent of the pool’s perimeter with woody vegetation. To quantify the degree of disturbance extending upland from water access points we laid out 3 parallel 2 m wide and 30 m long belt transects 10 m apart, centered at spring access points. Sampling locations were not shielded from potential donkey herbivory and disturbance by geographic barriers (e.g. cliffs). Along the entirety of each belt transect we counted the number of dung piles, and every 10 m we estimated % trampled ground, % undisturbed bare ground, % herbaceous cover, and % woody cover in 1 m^2^ quadrats (9 total per site). Although we collected data on plant cover by species, we did not include it in subsequent analyses because it was confounded by elevation and edaphic differences across sites, thus not directly capturing the effects of donkeys.

Instead of analyzing each of these response variables in sequential analyses, and because of their non-normality, we calculated Gower distance between sites based on all nine response variables and conducted a Principal Coordinates Analysis (PCoA), to identify the primary axes by which wetlands differed from each other. These first two axes (PCoA 1 and PcoA 2) explained 42.9% of total variation between sites (27.3 and 15.6% respectively). We then analyzed which factors explained differences between wetlands in this synthetic PCoA space using a PERMANOVA test in the R package ‘vegan’ v 2.5-6 [46] with 100,000 iterations. We included the presence of kills, the presence of cougars, elevation, and topographic complexity (S7 Fig) as independent variables.

To further test if these differences could be driven by underlying spatial gradients, we calculated a geographic distance matrix with the R package ‘geosphere’ v1.5-10[47]. We then conducted a Multiple Regression on Distances Matrices analysis with the function ‘MRM’ in the R package ‘ecodist’ v2.0.7 [48] with 1,000 iterations, which tested if the dissimilarity in wetland structure was explained by their distances from each other and thus underlying spatial gradients.

### Constraints on cougar predation

To understand the factors that constrained interactions between donkeys and cougars, we evaluated how landscape contexts influenced the probability of cougar presence at sites with feral donkeys. As others have shown [28], cougars require ambush cover provided either by vegetation or topography. We thus calculated a synthetic terrain-complexity variable from a 1/3 arc-second digital elevation model [49], which synthesized terrain roughness, terrain ruggedness, and slope with a Principal Components Analysis (PCA). PC1 explained 92.5% of total variation and was subsequently used as a terrain-complexity variable (S7 Fig).

To describe potential riparian ambush cover, we calculated total riparian area and the number of riparian patches by tracing the boundary of riparian vegetation in QGIS from satellite imagery (2016, Google), which was validated during field surveys. To test if anthropogenic landscape factors affected cougar presence, we recorded if springs were in the vicinity (within 500 m) of campsites or high-use recreation areas.

We then employed logistic generalized linear mixed effect models to understand the relative importance of campsites, terrain-complexity, total riparian area (on a log_10_ scale), and the presence of alternative prey at each site (presence or absence of bighorn sheep or mule deer) in predicting the presence of cougars and the presence of kills, as determined by field survey data (scat and sign) and camera-trap detections. Given that no kills occurred at campsites we excluded this factor from that analysis to prevent singularity failures. The region of each wetland was treated as a random block to control for spatial autocorrelation (S2 Fig, S1 Table 1). Alternative, more spatially explicit approaches, for instance using the R package ‘*spdep’* to create spatial neighborhood matrices and weights were not possible due to convergence failures in model. We excluded sites in two survey regions where cougar sign was absent across all sites, because the lack of sufficient sites within these block levels would weaken inferences [50] and because absence may have been driven by regional historic or stochastic factors.

Using multimodel selection techniques, we selected a final, most parsimonious model (based on AICc), which retained total riparian area and the presence of campsites as predictors for cougar presence, both of which were significant (*χ*^*2*^=6.5, p=0.01; *χ*^*2*^=4.7, p=0.03, S6 Fig). The most parsimonious model to explain the presence of kills included only terrain complexity, which was significant (*χ*^*2*^=5.0, p=0.03, S6 Fig).

## Acknowledgments

We would like to thank Death Valley National Park staff, particularly Alison Ainsworth and Mike Reynolds, and Mark Meyers of Peaceful Valley Donkey Rescue (PVDR) for assistance with field work logistics, equipment donations, and site information. Work was supported by the Australian Research Council (grant number DP180100272) and crowdfunding. We would like to thank Mark Davis and Martin Schlaepfer for helpful feedback on the manuscript.

## Notes

### Competing Interest Statement

The authors have declared no competing interest.

