## Supplemental Information for "Echoes of the late Pleistocene in a novel trophic cascade between cougars and feral donkeys"

**This PDF file includes:**

Supporting Text
Supporting Figures 1 to 9
Supporting Tables 1 to 3
Supporting References

### Supplemental Text

*Predator status in study areas*

Trail camera imagery was compiled from wetlands in the Sonoran and Mojave Deserts (Supplementary Table 1, Figure S2). The Sonoran Desert field sites occurred on Bureau of Land Management lands in central Arizona and data was collected from 2015-2018. Cougars are widely persecuted in this region, by recreational hunters and to protect livestock and to increase bighorn sheep production [1]. The predation event captured at one wetland (Fig. 2a-b) was by a young cougar whom we suspect was observed two years prior with their mother. Both of these cougars were suspected of hunting wild donkeys and wild horses at the site, as many of the equids (including adults) had injuries suggesting feline attacks and because of the rapid disappearance of all horse foals. The mother, who wore a GPS collar, appears to have since been killed by the state wildlife agency, which removes any cougar documented killing two bighorn sheep (*Ovis canadensis*) within a 6-month period [1].

The majority of fieldwork in the Mojave Desert occurred in Death Valley, California. Cougars are protected from recreational hunting and from most types of lethal management in the state of California. As the largest contiguous protected area in the continental United States, Death Valley may thus have the strongest protected population of cougars in the world.

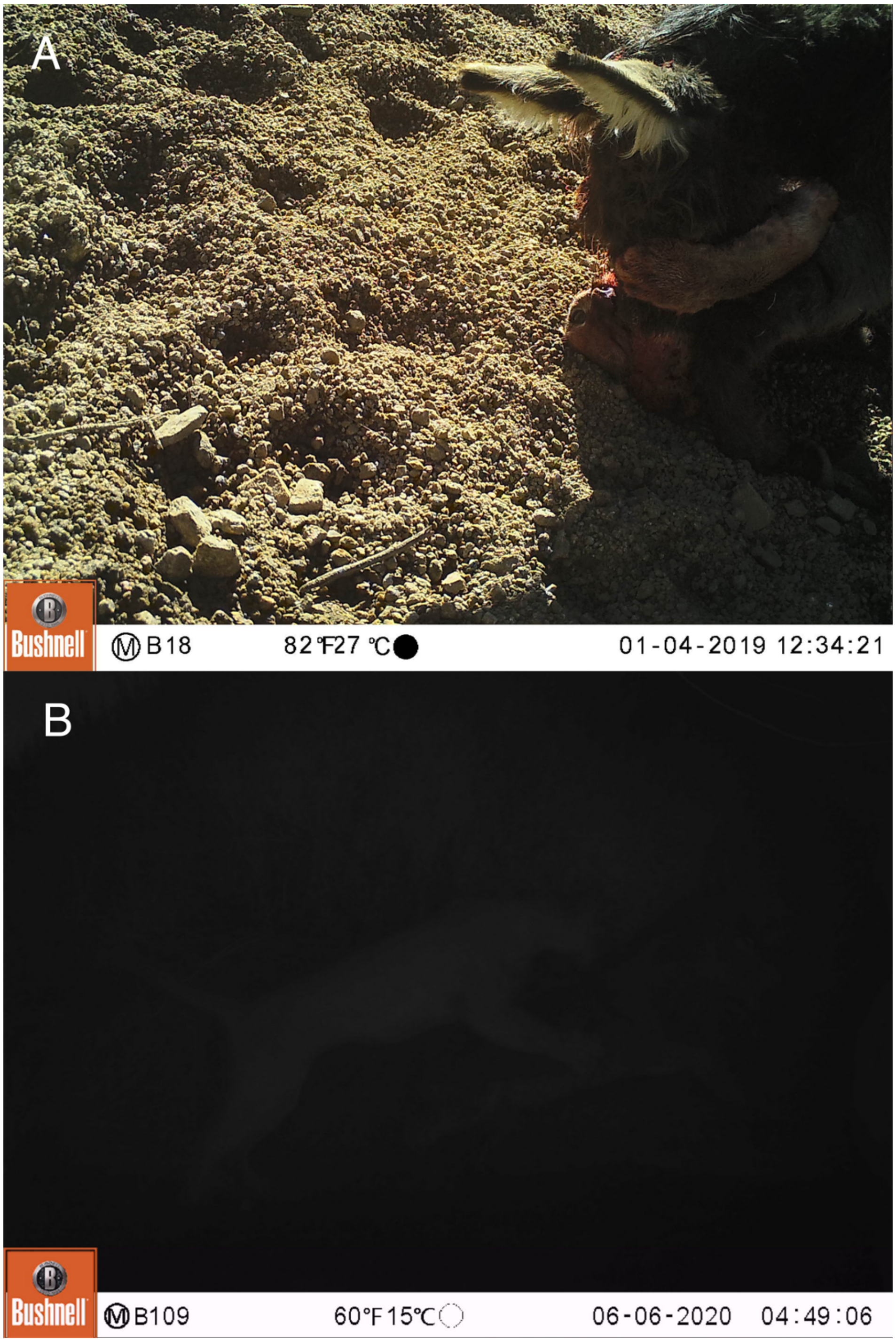


**S1 Fig. Original images from Fig. 2A and Fig. 2C, prior to tonal correction for underexposure.** Original files (.JPGs) are available upon request.

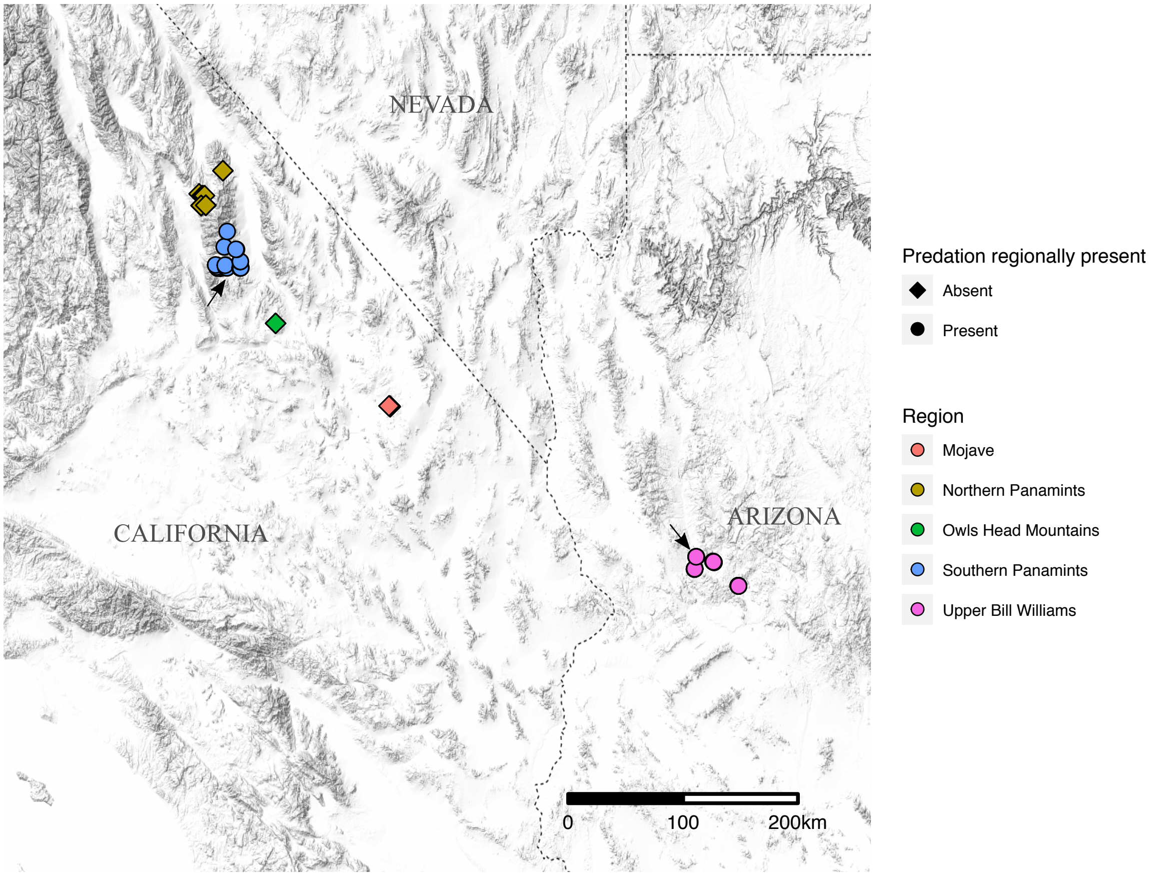


**S2 Fig. Study regions across the Sonoran ('Upper Bill Williams') and Mojave Deserts (all other study regions) of North America.** Color indicates region, with shape indicating whether predation was documented within the region. Black arrows point to sites where kills (Fig 2) were captured on camera traps. See Supplementary Table 1 for camera trap nights and details regarding which analyses included which sites.

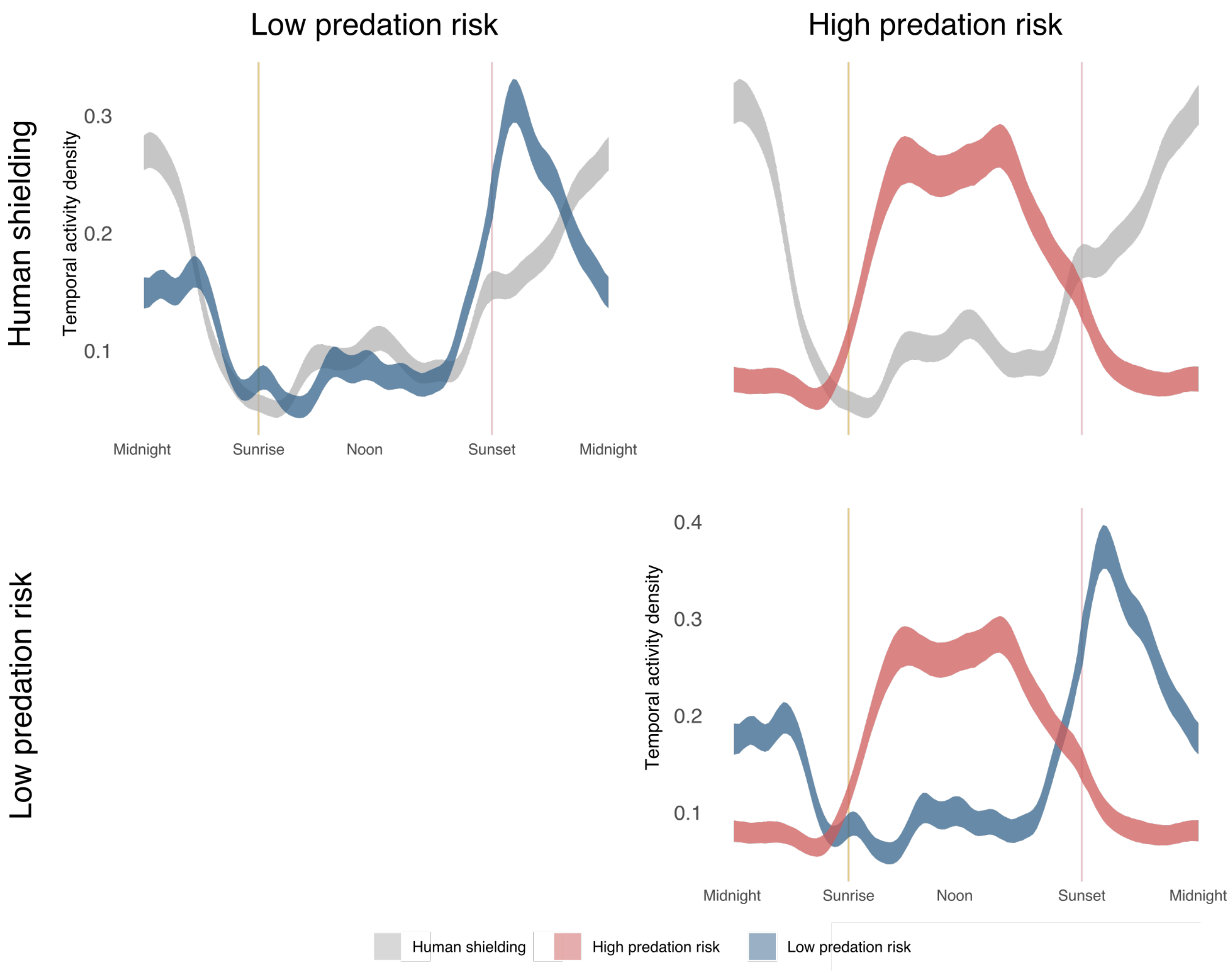


**S3 Fig. Pairwise overlap in temporal activity patterns (y axis) of donkeys under different levels of predation risk.** Ribbons indicate 95% confidence intervals across time (x-axis). Comparisons are as in a pairwise matrix, with rows and columns representing different predation risk categories.

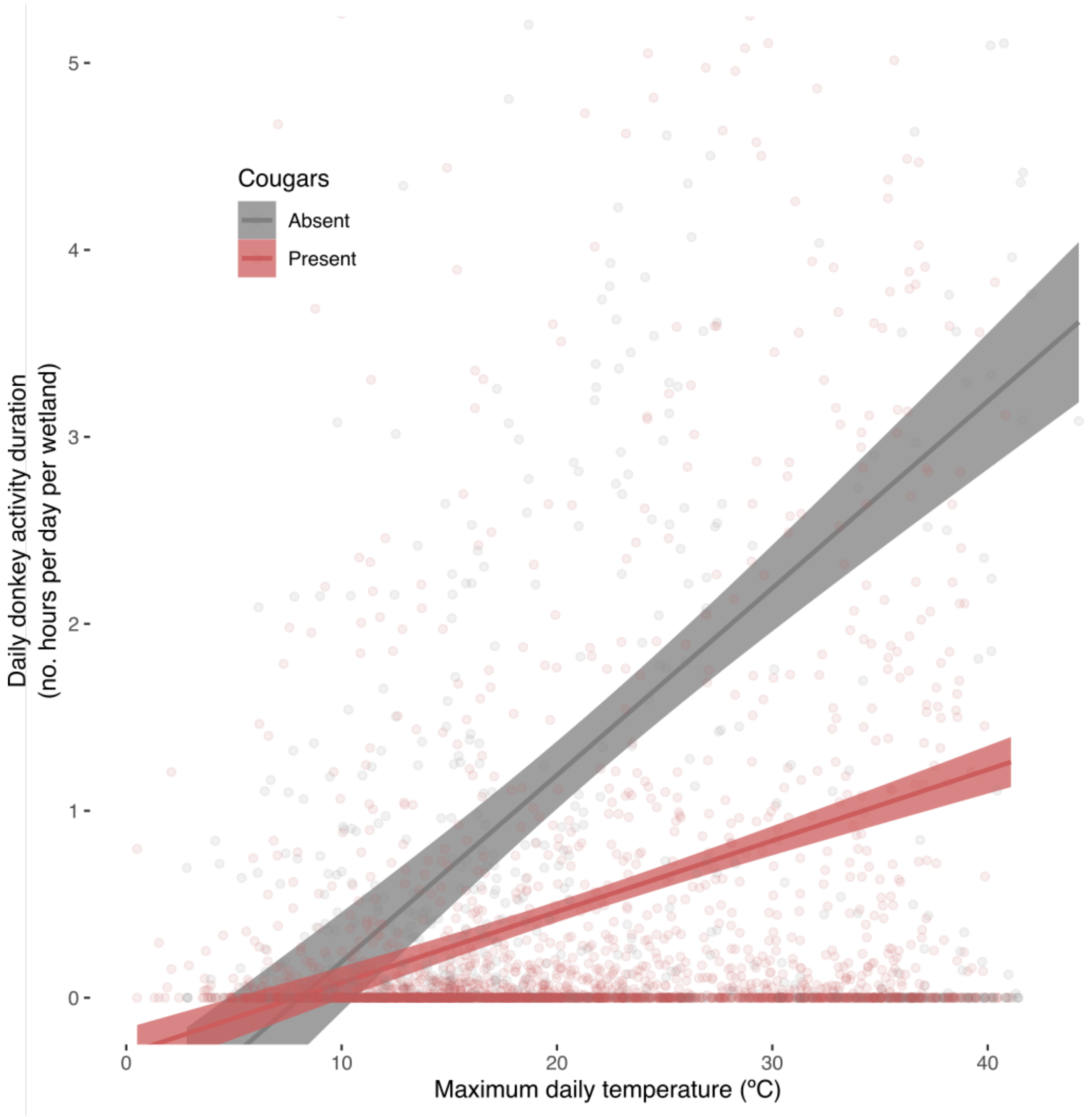


**S4 Fig. The presence or absence of cougars themselves affected daily activity of donkeys (hours/day/site), but to a lesser extent than the presence of predation**. Cougar presence was not included in the final most parsimonious model explaining daily occupancy. This suggests that behavioral responses are heightened at sites with high probability of successful ambush predation.

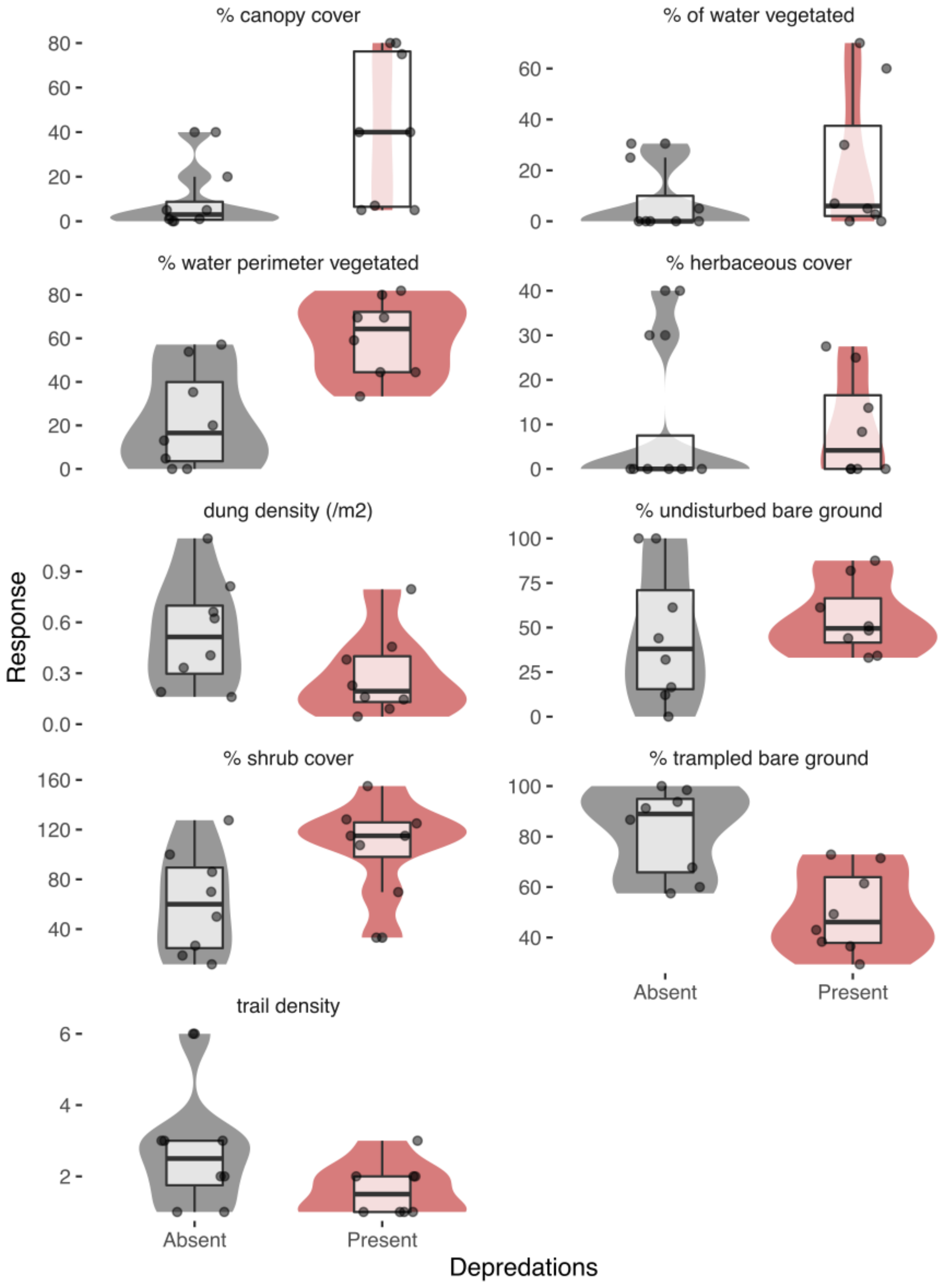


**S5 Fig. Individual soil and vegetation responses at desert wetlands, in areas with and without kills.** Fill indicates density distributions. Given their non-normality, these variables were analyzed synthetically with a PCoA and PERMANOVA test (Fig. 4).

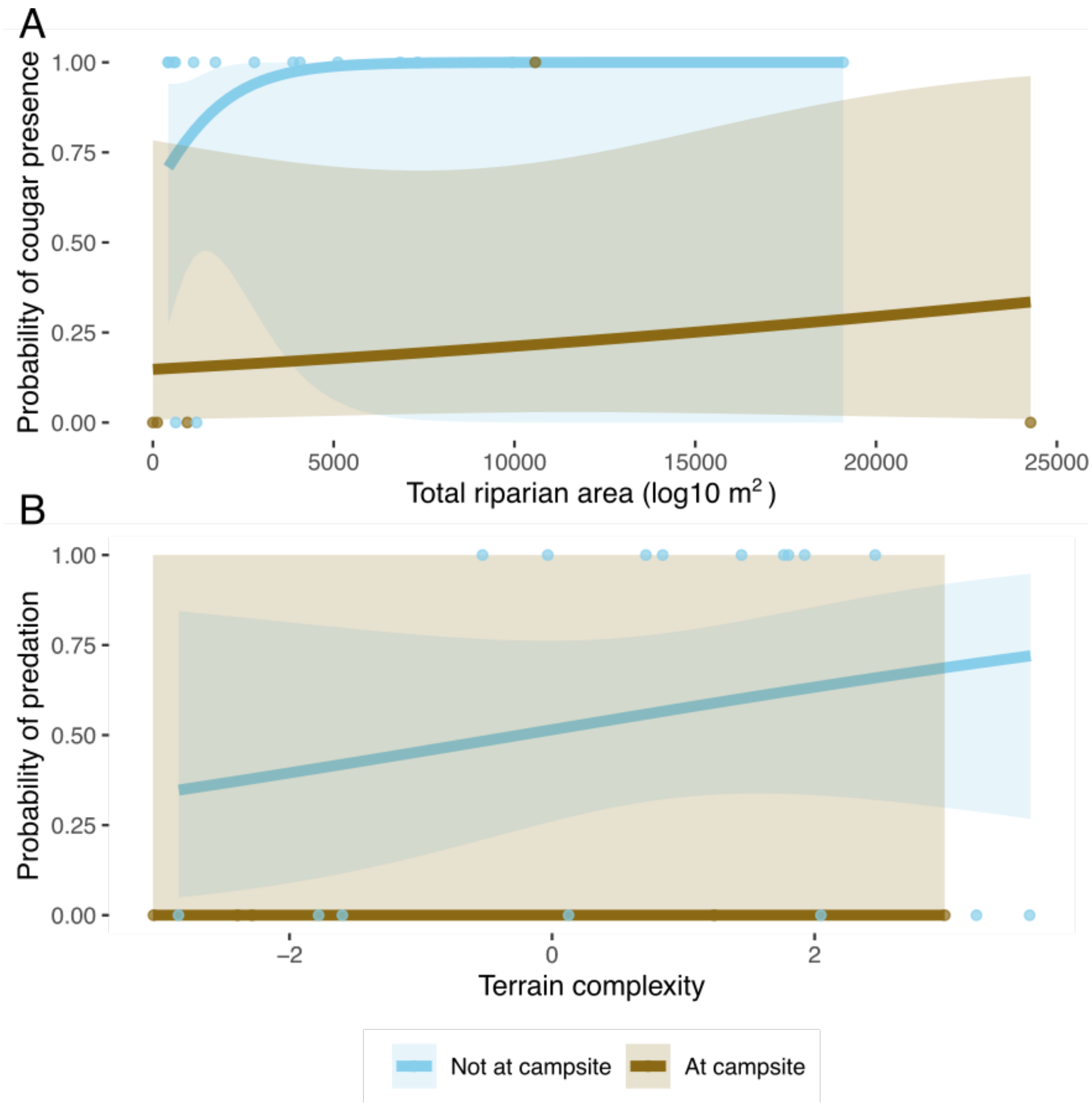


**S6 Fig. The most important factors influencing the presence of cougars and predation (e.g. kills) at desert wetlands in Death Valley National Park**. A. Riparian area and campsites were significant in predicting cougar presence (riparian area: $\chi^2$=6.5, p=0.01; campsites: $\chi^2$=4.7, p=0.03). B. The presence of kills was predicted by terrain complexity ($\chi^2$=5.0, p=0.03). Given that no kills occurred at human shielded campsite locations, that factor was excluded from analysis but is plotted here.

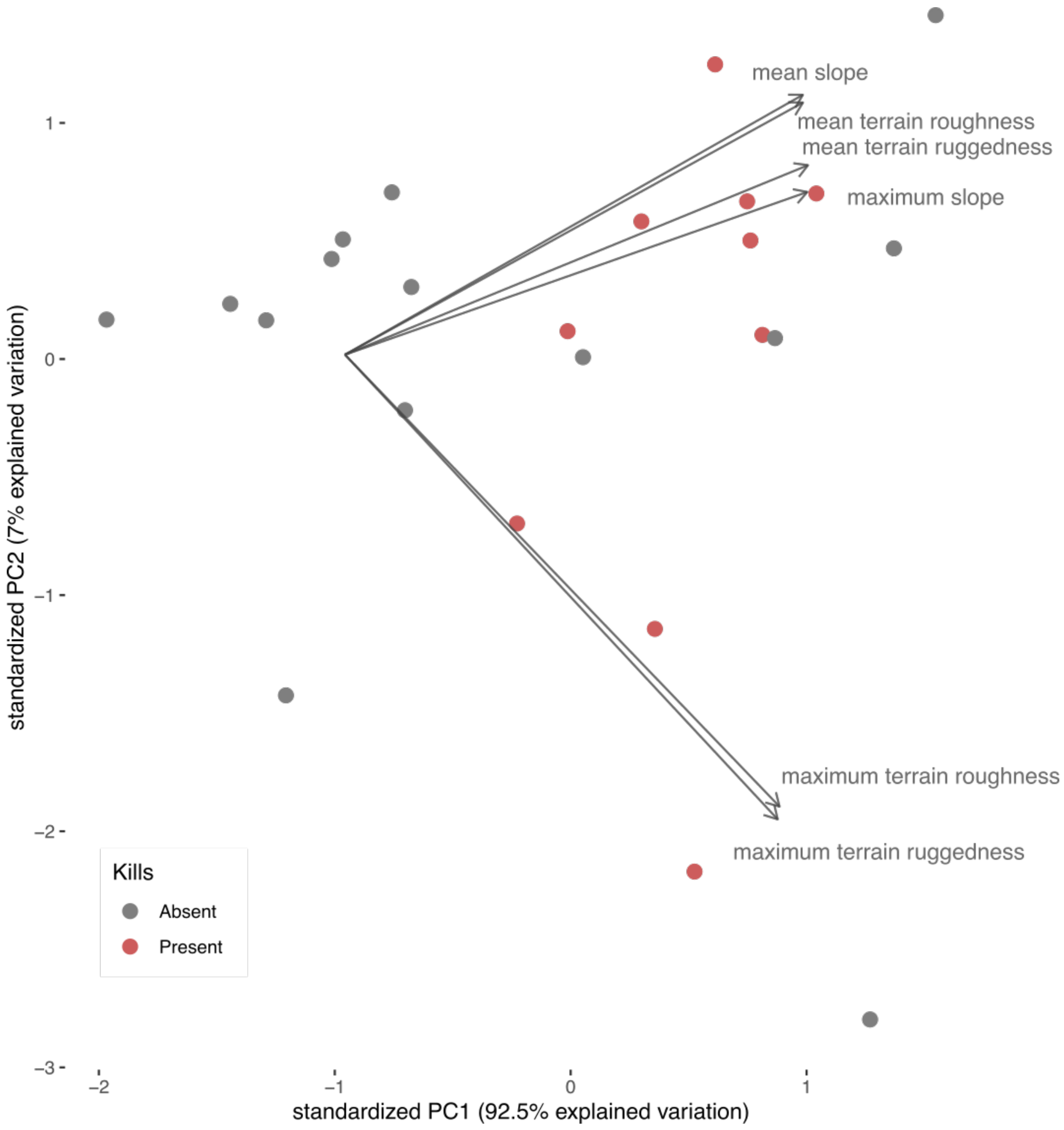


**S7 Fig. Terrain complexity Principal Components Analysis synthesizing an ensemble of terrain complexity metrics, including terrain roughness, slope, and ruggedness.** Terrain variables were calculated from a 1/3 arc-second digital elevation model (DEM) **[2]** and extracted within a 100m buffer for each site. PC1 (92.5% of total variation) was used in subsequent analyses as a synthetic estimate of terrain complexity to reduce model overfitting.

**S1 Table. Field site descriptions. Sites ranged across Sonoran and Mojave Deserts. Columns indicate region, corresponding to Supplementary Figure 1, and whether regional predation was documented**. Local predation details (e.g. at the wetland) include whether cougars were detected (through scat and/or camera surveys) and whether kills were detected (number in parentheses indicates number of kills). Risk category, used in temporal analyses (Fig. 3A) is indicated as is camera trapping summary with details on number of camera stations (e.g. individual cameras at a site), total trap nights across camera stations, and number of years of monitoring. To control for different regional metapopulation sizes, only the Southern Panamints were included in wetland occupancy analysis (Fig. 3B). Final column indicates whether sites were surveyed for donkey-related effects on wetland vegetation and soils.

| Wetland | Desert | Region | Regional predation | Local predation details | Risk category | Camera summary | Wetland effect survey |
|---|---|---|---|---|---|---|---|
| Anvil Spring | Mojave | Southern Panamints | yes | Cougars absent; kills absent | Human shielding | Cameras (1 station, 120 trap nights, 1 year) | surveyed |
| Arrastre Spring | Mojave | Southern Panamints | yes | Cougars present; kills present (1) | High predation risk | Cameras (2 stations, 86 trap nights, 2 years) | surveyed |
| Black Canyon | Sonoran | Upper Bill Williams | yes | Cougars present; Not surveyed | High predation risk | Cameras (4 stations, 432 trap nights, 2 years) | not surveyed |
| Blackwater Spring | Mojave | Northern Panamints | no | Cougars absent; kills absent | Low predation risk | No cameras | surveyed |
| Cane Springs | Mojave | Mojave | no | Cougars absent; kills absent | Low predation risk | Cameras (6 stations, 589 trap nights, 1 year) | not surveyed |
| Cattail Spring | Sonoran | Upper Bill Williams | yes | Cougars present; Not surveyed | High predation risk | Cameras (8 stations, 1088 trap nights, 1 year) | not surveyed |
| Five Mile Spring | Mojave | Southern Panamints | yes | Cougars present; kills present (2 + 1 kill captured on camera, Fig 2c-d) | High predation risk | Cameras (2 stations, 407 trap nights, 3 years) | surveyed |
| Greater View Spring | Mojave | Southern Panamints | yes | Cougars absent; kills absent | Human shielding | Cameras (1 station, 171 trap nights, 2 years) | surveyed |

| | | | | | | | |
|---|---|---|---|---|---|---|---|
| Greenwood Spring | Sonoran | Upper Bill Williams | yes | Cougars present; kills present (Kill captured on camera Fig 2a-b) | High predation risk | Cameras (6 stations, 950 trap nights, 2 years) | not surveyed |
| Hackberry Wash | Sonoran | Upper Bill Williams | yes | Cougars absent; Not surveyed | High predation risk | Cameras (6 stations, 501 trap nights, 2 years) | not surveyed |
| Hatchet Spring | Mojave | Southern Panamints | yes | Cougars present; kills absent | High predation risk | No cameras | surveyed |
| Hidden Valley Spring | Mojave | Southern Panamints | yes | Cougars present; kills absent | High predation risk | Cameras (1 station, 57 trap nights, 2 years) | surveyed |
| Little Spring | Mojave | Southern Panamints | yes | Cougars present; kills absent | High predation risk | Cameras (2 stations, 533 trap nights, 3 years) | surveyed |
| Lost Spring | Mojave | Southern Panamints | yes | Cougars present; kills present (1) | High predation risk | Cameras (5 stations, 1108 trap nights, 2 years) | not surveyed |
| Lower Galena | Mojave | Southern Panamints | yes | Cougars present; kills present (3) | High predation risk | Cameras (1 station, 253 trap nights, 2 years) | surveyed |
| Lower Tuber Spring | Mojave | Northern Panamints | no | Cougars present; kills absent | Low predation risk | Cameras (3 stations, 318 trap nights, 2 years) | surveyed |
| Mesquite Spring | Mojave | Southern Panamints | yes | Cougars absent; kills present (1) | High predation risk | Cameras (3 stations, 430 trap nights, 2 years) | surveyed |
| Mud Spring | Mojave | Northern Panamints | no | Cougars absent; kills absent | Low predation risk | Cameras (1 station, 184 trap nights, 3 years) | surveyed |
| N Fork Johnson Canyon | Mojave | Southern Panamints | yes | Cougars present; kills present (1) | High predation risk | No cameras | surveyed |
| Owls Head Spring | Mojave | Owls Head Mountains | no | Cougars absent; kills absent | Human shielding | Cameras (2 stations, 199 trap nights, 2 years) | surveyed |
| Powerful Women Spring | Mojave | Southern Panamints | yes | Cougars present; | High predation risk | Cameras (1 station, 444 trap nights, 2 years) | surveyed |

| | | | | | | | |
|---|---|---|---|---|---|---|---|
| | | | | kills present (1) | | | |
| Upper Galena | Mojave | Southern Panamints | yes | Cougars present; kills present (3) | High predation risk | Cameras (1 station, 266 trap nights, 2 years) | surveyed |
| Upper Tuber | Mojave | Northern Panamints | no | Cougars present; kills absent | Low predation risk | Cameras (3 stations, 318 trap nights, 1 year) | not surveyed |
| Warm Springs | Mojave | Southern Panamints | yes | Cougars present; kills present | High predation risk | No cameras | not surveyed |
| Wildrose | Mojave | Northern Panamints | no | Cougars absent; kills absent | Human shielding | Cameras (1 station, 251 trap nights, 2 years) | not surveyed |
| Willow Spring | Mojave | Southern Panamints | yes | Cougars present; kills present (10) | High predation risk | Cameras (4 stations, 1119 trap nights, 3 years) | surveyed |

**S2 Table. Full PERMANOVA model results**. PERMANOVA was conducted with the function ‘adonis2’ in the R package ‘vegan’ v2.5-6 **[3]** with 100,000 iterations.

| Variable | Degrees of freedom | Sum of squares | $R^2$ | F | p-value |
|---|---|---|---|---|---|
| kills present | 1 | 0.31 | 0.20 | 3.54 | 0.00014 |
| cougars present | 1 | 0.10 | 0.06 | 1.11 | 0.33 |
| terrain complexity | 1 | 0.10 | 0.06 | 1.13 | 0.31 |
| elevation | 1 | 0.10 | 0.07 | 1.19 | 0.27 |
| Residual | 11 | 0.95 | 0.61 | | |
| Total | 15 | 1.56 | 1 | | |

**S3 Table. Evidence of predation on introduced wild equids by extant predators.** The possibility that predators can influence introduced equids is frequently dismissed in policy and research. However, growing evidence suggests that extant predators have a greater capacity to influence equids than usually considered. In Africa and Asia, *Panthera uncia* [4], *Panthera pardus* [5], *Panthera leo* [6], and *Crocuta crocuta* [7] have been documented predating domestic or wild (non-introduced) equids and are displayed in Fig. 6, but are not included here as they do not overlap with introduced equid populations.

| Predator | Description |
|---|---|
| Cougar (*Puma concolor*) | *Horses*<br>Cougars have been documented preying on horses, though studies often have mixed results regarding the importance of horses in cougar diets [8, 9]. Predation on foals [10] has been shown to have significant effects on population growth in some study areas [11-13]. However, a 25-year study on a heavily predated and unmanaged horse population found that horses began avoiding their historic summer rangelands (higher elevation with more vegetative cover), presumably to avoid cougar predation [14].<br><br>While predation on juveniles has been most commonly documented [11-14], Andreasen [15] documented predation on adult horses as well (estimated ~420kg in her study area) by both male and (smaller-bodied) female cougars. In some study populations, horses were the primary prey item, particularly in mountain ranges with high densities of horses.<br><br>Cougar influences on horse populations are likely sensitive to even low levels of persecution. For example, cougar predation appeared to drive population decline among horses in the Pryor Mountains of Montana and Wyoming until the removal of 3 cougars by humans. Following this, horse population growth rapidly increased [16].<br><br>*Donkeys*<br>Cougar predation on donkeys had not been documented in the literature until this study. |
| Gray wolf (*Canis lupus*) | *Horses*<br>Horses can be a major component of wolf diets in some regions, including in Europe [17-19] and in Canada, where wolves and horses overlap [20, 21]. However, wolves do not currently overlap with most introduced horse populations in North America (Figure 6) given historic extirpation of wolf populations from much of the United States and Mexico.<br><br>The cursorial and pack hunting strategy of wolves suggests that they would be able to hunt introduced horses and donkeys in areas without sufficient ambush cover for successful cougar predation.<br><br>*Donkeys*<br>Wolf predation has been documented on domestic donkeys in Iberia [22] and gray wolves have been observed chasing reintroduced Asiatic ass (*Equus hemionus*) in Israel (Gavin Bonsen *personal communication*).<br><br>Wolves and donkeys do not currently overlap in North America (Figure 6). Whether gray wolves could reestablish in the desert environments occupied by donkeys remains unknown, yet is plausible given the presence of gray wolves in hot deserts in Eurasia [23] and anecdotal records of gray wolves in hot, hyper-arid parts of the Sonoran Desert, such as the Pinacate Biosphere of Northern Mexico [24]. |
| Jaguar (*Panthera onca*) | *Horses and donkeys*<br>Jaguars can be major predators of domestic livestock, including horses and donkeys [25, 26]. To the best of our knowledge, no study has been conducted on how jaguars may influence feral equids in South America or Mexico, where they may overlap. Jaguars have experienced significant range contractions in North and South America [23]. Their reestablishment, |

| | |
|---|---|
| | particularly in the Southwestern United States, could lead to increased predation pressure on adult equids. |
| Dingo (*Canis dingo*) | *Horses*<br>Horses are commonly recorded in dingo scats, albeit at low frequencies [27]. Dingo predation on a feral horse foal was observed in the Painted Desert of Australia (ADW, *personal observation*) and dingo packs are known to kill feral horses in the Snowy Mountains, including foals at least as old as 6 months in age[28]. This has been thought to perhaps explain the larger size of dingo packs in that region [29]. Experimental work found that dingo howls did not elicit maternal protectiveness responses among feral horse mares but that dominant stallions responded by spending more time in close proximity to foals, suggesting that dominant stallions may play an important role defending foals from dingo predation[30].<br><br>*Donkeys*<br>Donkeys have been recorded in dingo scats and dingo packs have been observed in pursuit of donkeys (Chris Henggeler *personal communication* Aug. 2019). Furthermore, the protection and stabilization of dingo populations has been strongly linked to reduced donkey abundance [31]. However, dingo predation on donkeys has not been directly recorded. |
| Brown bear (*Ursus arctos*) | *Horses*<br>Brown bears have been documented predating domestic horses in Spain, including both yearlings and adults [32]. In Alberta, Canada, grizzly bear (*Ursus arctos horriblis*) have been observed chasing horses and are suspected to be a major cause of mortality (Paul Boyce, *personal communication*).<br><br>*Donkeys*<br>Given widespread range contractions, brown bears and introduced wild donkeys do not currently cooccur (Figure 6). Gobi desert grizzly bears co-occur with Asiatic ass (*Equus hemionus*), but whether they predate these donkey-like equids is unknown [33]. If grizzly bears were to reestablish in arid regions of North America [23] then interactions with wild donkeys could be possible. |
